# Synchronization of Hes1 oscillations coordinate and refine condensation formation and patterning of the avian limb skeleton

**DOI:** 10.1101/157446

**Authors:** Ramray Bhat, Tilmann Glimm, Marta Linde-Medina, Cheng Cui, Stuart A. Newman

## Abstract

The tetrapod appendicular skeleton is initiated as spatially patterned mesenchymal condensations. The size and spacing of these condensations in avian limb buds are mediated by a reaction-diffusion-adhesion network consisting of galectins Gal-1A, Gal-8 and their cell surface receptors. In cell cultures, the appearance of condensations is synchronized across distances greater than the characteristic wavelength of their spatial pattern. We explored the possible role of observed oscillations of the transcriptional co-regulator Hes1 in this phenomenon. Treatment of micromass cultures with DAPT, a γ-secretase inhibitor, damped Hes1 oscillations, elevated Gal-1A and -8 mRNA levels, and led to irregularly-sized and fused condensations. In developing limb buds, DAPT led to spatially non-uniform Hes1 expression and fused and misshapen digits. Periodicity in adhesive response to Gal-1A, a plausible Hes1-dependent function, was added to a previously tested mathematical model for condensation patterning by the two-galectin network. The enhanced model predicted regularization of patterning due to synchronization of Hes1 oscillations and resulting spatiotemporal coordination of its expression. The model also predicted changes in galectin expression and patterning in response to suppression of Hes1 expression, which were confirmed in in vitro experiments. Our results indicate that the two-galectin patterning network is regulated by Hes1 dynamics, the synchronization of which refines and regularizes limb skeletogenesis.

## Introduction

The appendicular skeleton in tetrapods is characterized by a stereotypical pattern that consists of an increase in number of bony elements arranged in series, along the proximodistal (body-to-digit tip) axis. Each bone is prefigured by a cartilaginous element that in turn differentiates from a cellular condensation: a tight aggregate of somatopleure-derived mesenchymal cells (Newman and Bhat, 2007; Newman et al., 2018). The digits in the tetrapod limb (collectively called the autopod) derive from such condensations, which arise spaced well-apart from each other. Investigations into the molecular mechanism(s) that pattern the pre-digit condensations, i.e., determinants of their size, number and spatial separation, have implicated a variety of molecules, including cell-cell adhesion proteins, extracellular matrix and matricellular proteins and their interacting partners, and diffusible morphogens (reviewed in (Newman and Bhat, 2007) and (Newman et al., 2018)). Of these, galectin-1A (Gal-1A) and galectin-8 (Gal-8) are, along with their putative receptors, the earliest proteins found to be expressed in a condensation-specific fashion within developing chicken limb mesenchyme (Bhat et al., 2011). They are also functionally involved in the patterning process: together they constitute an activator-inhibitor-adhesion reaction-diffusion network with well-characterized dynamics that determines the sizes and spacing of precartilage cell condensations in vitro and of digits in ovo (Bhat et al., 2011; Glimm et al., 2014).

Morphogenetic processes specified via reaction-diffusion-based mechanisms are constrained in the spatial scale of their activity by the local nature of diffusing interactors. For such systems to form reliable patterns over distances greater than their characteristic wavelengths (e.g., across the developing digital plate or a micromass culture) they must incorporate one or more features that promote global coordination. The spatiotemporal synchronization of cell fates during development has been actively investigated over the last two decades in the context of vertebrate segmentation. Specifically, synchronized oscillations in the expression of genes belonging to signaling pathways such as Notch and Wnt constitute a molecular clock that appears to determine the periodicity of the sequential formation of the somites that give rise to vertebrae and other tissues of the body (Aulehla and Pourquie, 2006). The frequency of the oscillations in expression of *hes1*, a gene encoding a transcription factor downstream of Notch in the unsegmented presomitic mesoderm, correlates with the formation of somites (Palmeirim et al., 1997). The chicken homolog of *hes1* (also known as *c-hairy1*), is also expressed within distal undifferentiated limb mesenchyme and its overexpression leads to shortening of skeletal elements (Vasiliauskas et al., 2003). The Notch pathway appears to be active within high-density micromass cultures of precartilage limb mesenchyme and its pharmacological inhibition led to increased size and irregular shapes of condensations and accelerated their formation (Fujimaki et al., 2006). In particular, mice in which both presenilin genes (principal constituents of the γ-secretase complex essential for juxtacrine signaling), or Jagged 2 were deleted show patterning defects that include impairment in interdigital apoptosis and distal syndactyly (Jiang et al., 1998; Pan et al., 2005).

Here we present experimental evidence that signaling mediated by the Hes1 transcriptional modulator is intrinsic to the two-galectin patterning network for precartilage condensation. We show furthermore, that the sustained spatially correlated oscillatory expression of the *hes1* gene is involved in the synchronized formation of condensations in vitro and the coordinated emergence of patterned digits in vivo. We incorporate the effects of oscillatory dynamics on galectin expression in our mathematical model to provide a mechanistic understanding of how the entrainment of oscillations helps coordinate the spatial emergence of the condensation pattern and sharpen the boundaries of forming skeletal elements. Our simulations, in conjunction with our experimental results and those in the literature suggest that the effect of synchronization of oscillations is to make Hes1 status uniform across the tissue mass so that globally organized patterning signals evoke corresponding effects at distant sites.

## Results

### Limb precartilage mesenchymal cells condense simultaneously over millimeter distances in culture

When the digital plate mesenchyme of embryonic chicken limb buds is dissociated and cultured at a high density, cells begin aggregating within a few hours, forming regularly spaced foci (“protocondensations”) marked by the expression of the two galectins Gal-1A and Gal-8 (Bhat et al., 2011). These foci then mature into classically described precartilage condensations (Frenz et al., 1989; Paulsen and Solursh, 1988). We tracked the formation of the condensations in time across high-density micromass culture (∼3 mm diameter) with ∼5% of the limb mesenchymal cells expressing H2B-GFP. Morphological condensations, first evident by fluorescence optics by about 48 h after the culture was initiated (as in similar cultures viewed by phase contrast microscopy), appeared nearly simultaneously across the entire micromass (Fig. 1, Movie S1). We have shown previously that the morphogenesis of individual condensations and the inhibition of formation of others in their immediate vicinity is dependent on local regulatory interactions between two galectins and their counterreceptors (Bhat et al., 2011; see also Lorda-Diez et al. (2011)). Here, we find in addition that the morphogenetic events mediated by this network are synchronized across a non-local “morphogenetic field” (Gilbert and Sarkar, 2000; Levin, 2012), represented by the culture micromass and the digital plate.

**Figure 1:**
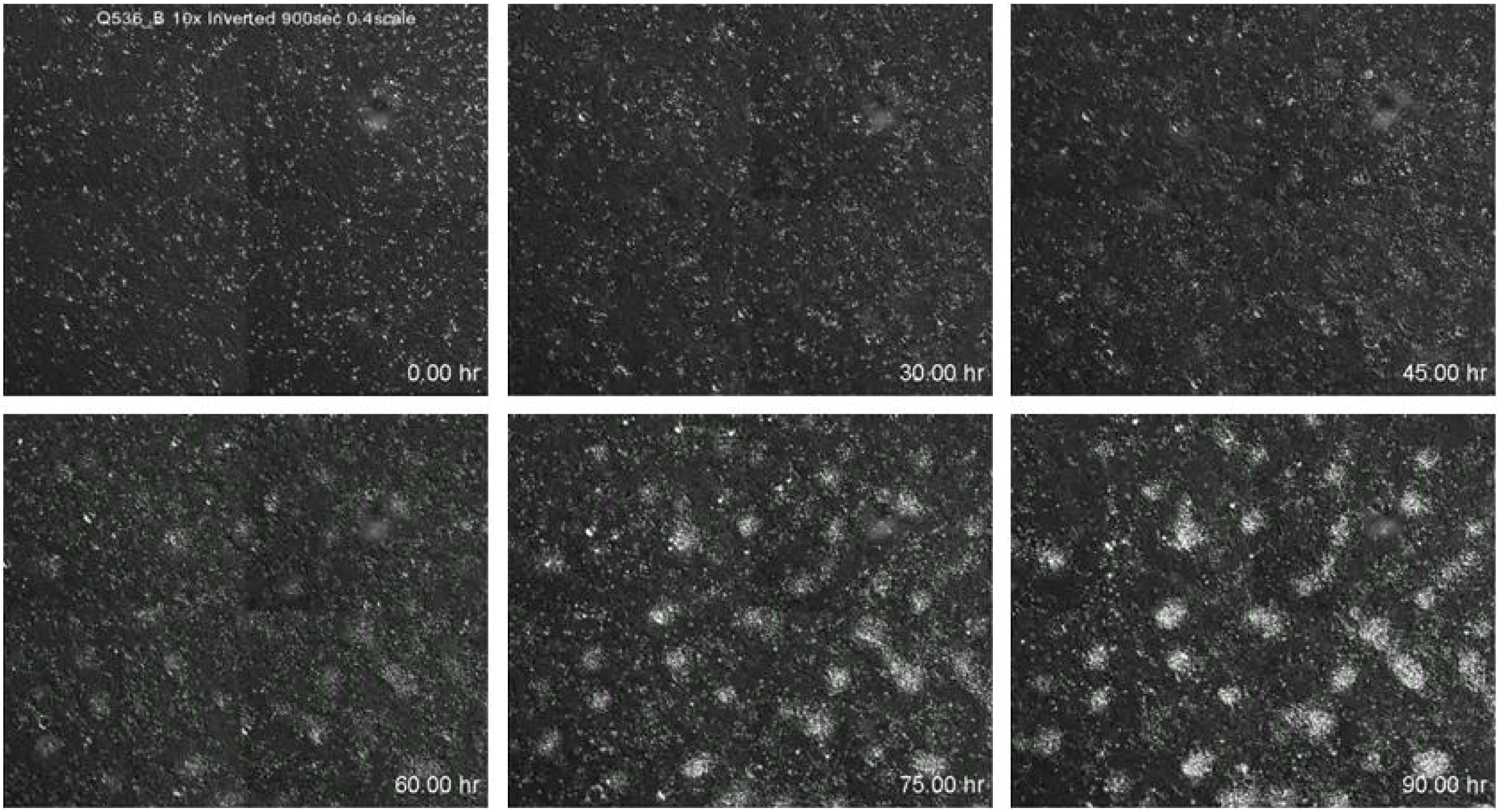
In vitro precartilage mesenchymal condensations emerge in spatiotemporal coordination. Time lapse fluorescent micrographs of a high density micromass culture of precartilage limb mesenchymal cells with ∼5% cells expressing H2B-GFP. Condensations of mesenchymal cells appear simultaneously across the field of view suggesting that condensation morphogenesis is coordinated across the micromass. This figure is derived from a time lapse video (Movie S1) that was acquired once. The coordinate appearance of condensations was confirmed in more than a dozen micromass cultures sampled by still Hoffman Modulation Contrast photomicrography over the same duration

### Expression of *hes1* undergoes oscillation during condensation patterning in vitro

The evidence of temporal synchrony in condensation morphogenesis was suggestive of other cases of temporally coordinated pattern formation during animal development. During vertebrate somitogenesis, signaling in the presomitic mesoderm synchronizes *hes1* oscillations across the width of the presomitic mesoderm, constituting each segmenting band of tissue as a coherently acting tissue mass, i.e., morphogenetic field (Palmeirim et al., 1997). To test whether Hes1 oscillations play a similar role in limb skeletal patterning, we assayed the expression of its encoding gene *hes1* in cultures of precartilage leg mesenchyme using real-time PCR. This permitted a quantitative estimation of temporal expression dynamics in a population of moving cells. When assayed at regular time points (with intervals ranging from 30 min to 3 h), between 0 h and 33 h post-incubation (the time window in which protocondensations, and then condensations appear), the expression of *hes1* was found to rise and fall periodically between 12 h and 24 h (Fig. 2Ai). When the interval between assay time points was decreased to 1 h or ½ h (to detect oscillations of potentially lower periodicities), the expression plots nevertheless showed temporal fluctuations that were diagnostic of larger oscillatory time periods (Fig. 2Aii,iii). Interestingly, the expression of *hes1* before 12 h and after 24 h post-plating was non-oscillatory (Fig. 2Ai), suggesting that the oscillation of Hes1 mRNA was tightly associated temporally with the period of condensation patterning and refinement. Using the Lomb-Scargle algorithm, devised to quantify periodicity of a signal (Glynn et al., 2006), the expression of *hes1* was found to oscillate with a periodicity of 6 h (Fig. 2B). We assayed the protein levels of Hes1 through the protocondensation period by indirect immunocytochemistry, finding that diffuse staining for Hes1 in early micromasses at 9 h gave way to a more nodular condensation-specific staining for Hes1 at 24 h (Fig. 2C).

**Figure 2:**
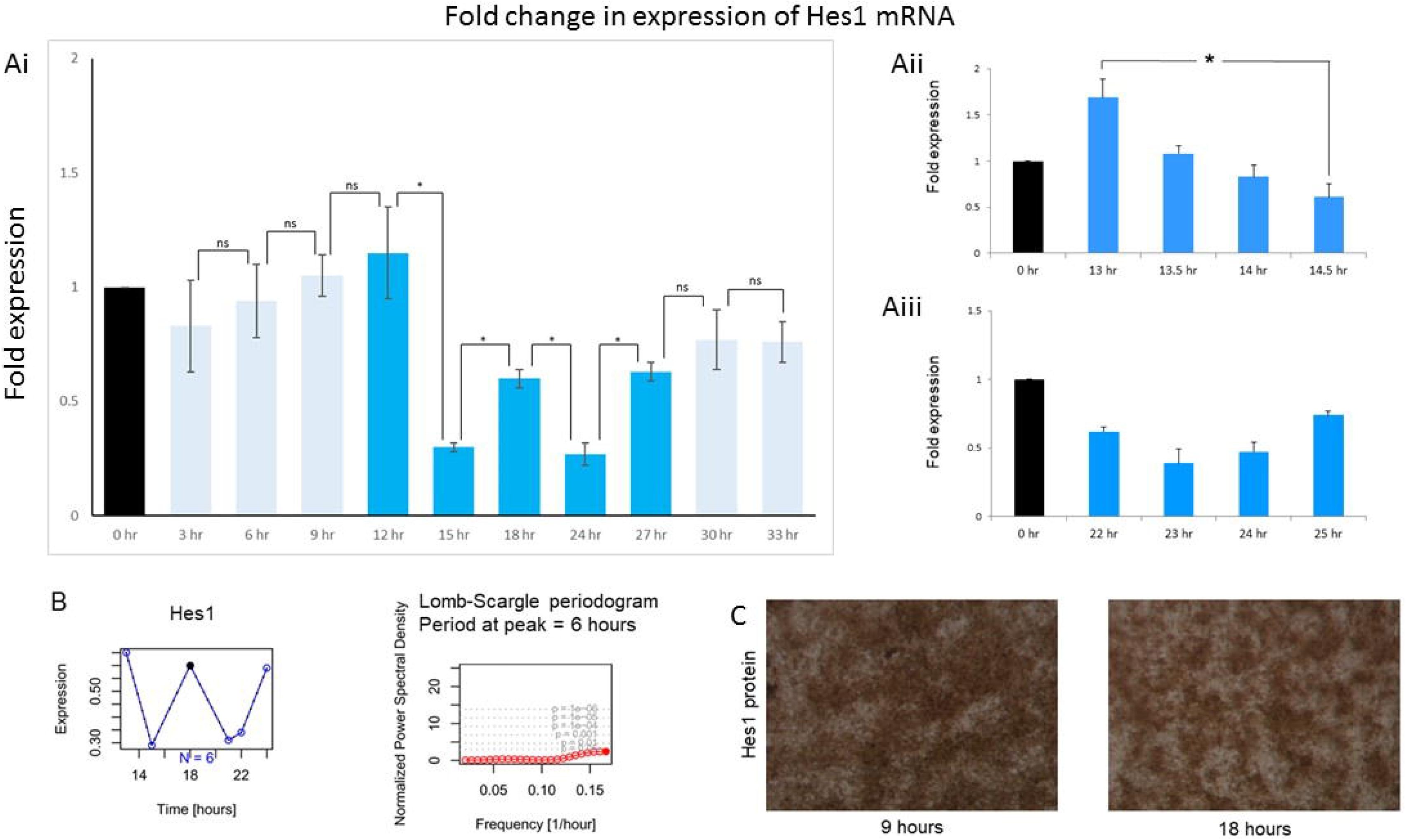
Gene expression of Hes1 undergoes oscillation in condensing mesenchymal cells within high density micromasses. (Ai) Assay for Hes1 gene expression in developing limb bud micromass cells using real-time quantitative PCR shows an oscillatory temporal pattern between 12 and 24 hours after induction of culture with no oscillatory signatures before or after this time window. (Aii and iii) Hes1 expression in cell cultures measured every half hour (above) and one hour (below) using real-time quantitative PCR show temporal dynamics that were diagnostic of the oscillatory expression seen in Ai. (B) Analysis of Hes1 expression periodograms assayed >3 times with time intervals of 3 h as well as with time intervals of 1 h and 0.5 h, when integrated using Lomb-Scargle algorithm reveals a periodicity of 6 hours for oscillation of Hes1 mRNA levels. (C) Localization of Hes1 protein in micromass culture, assayed through indirect immunocytochemistry shows a more diffuse staining at 9 h (prior to condensation formation) settling into a more condensation-scale pattern at 18 h. Error bars represent S.E.M. *p < 0.05.

### Inhibition of γ-secretase alters condensation patterning dynamics in vitro and leads to disruption in digit morphologies in ovo

In order to test whether suppression of Hes1 activity alters condensation patterning, we treated limb micromass cultures with DAPT. DAPT inhibits the cytoplasmic cleavage of the Notch intracytoplasmic domain but also can potentially affect the function of other cell signaling pathways in which presenilin-1-dependent intramembrane cleavage operates, including those dependent on TGF-β (Blair et al., 2011) (Hemming et al., 2008) and Wnt (Mi and Johnson, 2007)) signaling. In fact, both TGF-β- and Wnt-based signaling have been shown to regulate the transcription of Hes1 in a Notch-independent manner (Xing et al., 2010) (Peignon et al., 2011). In DAPT-treated micromass cultures, there was a sustained decrease in the expression of the gene encoding Hes1, with no evidence of the oscillatory dynamics seen in control (compare Fig. 3A to Fig. 2A). We analyzed the changes in condensation patterning associated with inhibition of γ-secretase by focusing on condensation number, condensation size and inter-condensation distance. Condensation number was decreased by DAPT, which could be accounted for by the fusion of neighboring condensations (Fig. 3B). Frequency distributions of condensation sizes showed both increased smaller and larger condensations (Fig. 3C left), whereas the distribution of intercondensation distance showed a left shift (Fig. 3C right), indicating irregular patterns of condensations upon DAPT treatment.

**Figure 3:**
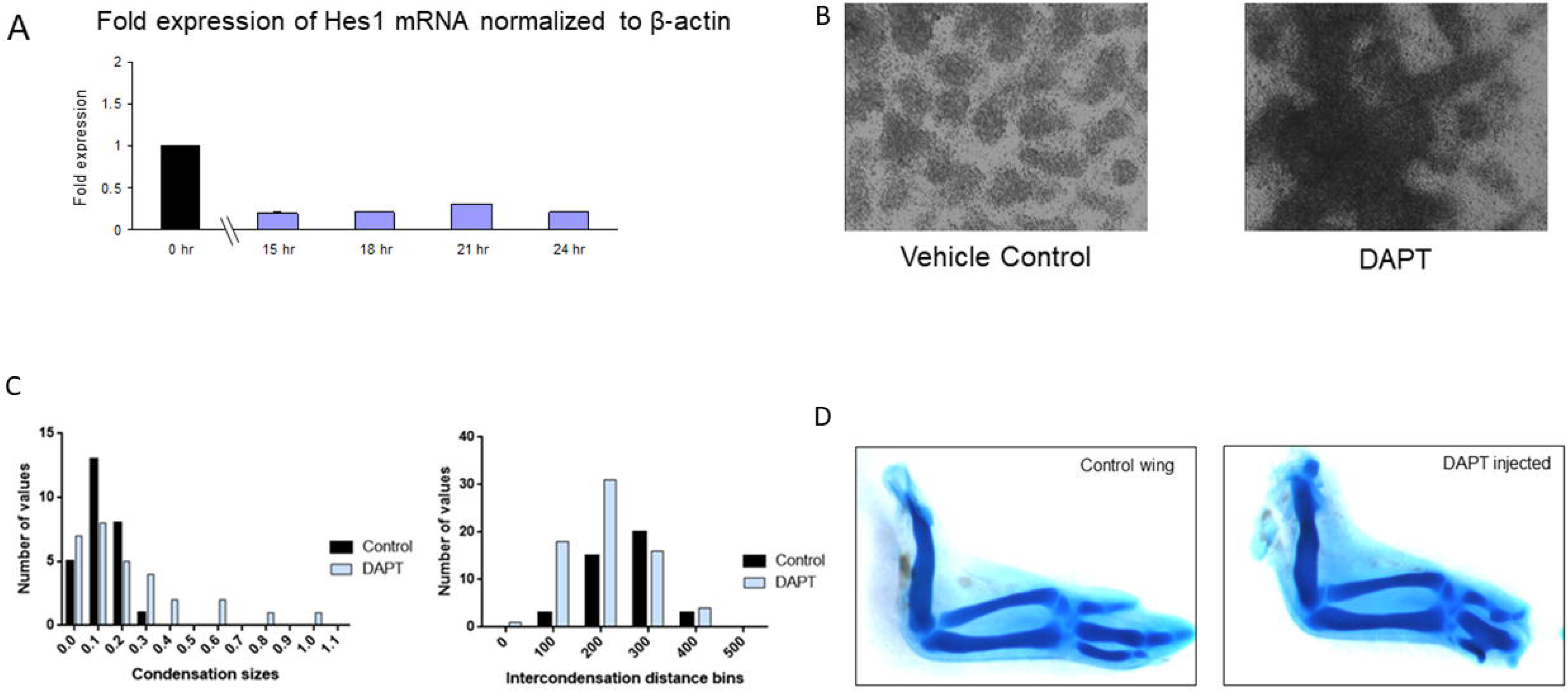
The γ-secretase inhibitor DAPT enhances the gene expression of Gal-1A and Gal-8. (A) Treatment of limb micromass cultures with DAPT results in a damping in the expression of Hes1 mRNA when assessed between 15-24 hours post initiation of culture, using RTqPCR. (B) Treatment of precartilage limb micromasses with □-secretase inhibitor DAPT increased fusion of condensations with a resultant significant change in intercondensation distance and mean condensation size (p<0.05). (C) Frequency distributions of condensation sizes and numbers in the absence and presence of DAPT. (D) In ovo, injection of DAPT into developing wing autopodium resulted in alteration in digit morphologies with fusions between digital elements.

Our in vitro experiments suggested that γ-secretase-dependent Hes1 oscillations act to coordinate spatial patterning of condensations. We next sought to investigate if such an effect was also evidenced *in ovo*. Potential pattern refinement dependent on oscillator synchronization requires physical continuity throughout the morphogenetic field. We introduced tantalum foil barriers into prospective autopodial mesenchyme of avian wing buds and assayed for spatial expression of *hes1* by in situ hybridization. For each wing bud tested, the expression pattern was altered relative to the pattern in the contralateral counterpart (Fig. S1). The foil barriers typically led to disparate staining patterns on its two sides, where the contralateral limb showed uniform high or low expression or a symmetrical expression pattern.

Earlier reports indicated that insertion of foil barriers at precondensation stages of autopodial development alters digit patterning (Rowe and Fallon, 1982). By our hypothesis this effect could be mediated by the disturbance of *hes1* expression synchrony resulting from this mechanical intervention. We used an alternative, biochemical means of perturbing *hes1* expression synchrony by injecting DAPT into the prospective autopod of developing wing buds. DAPT injection altered the spatial pattern of Hes1 mRNA expression as assayed by in situ hybridization: a distinct zone of inhibition in Hes1 expression was observed followed by a new locus of Hes1 expression emerging in the distalmost mesenchymal cells subjacent to the AER. No such locus of expression was seen in the contralateral vehicle-injected wing buds (Fig. S2). When evaluated post-chondrogenesis, this treatment was seen to result in fusion between phalanges and other abnormalities in digit and proximal element morphologies (Fig. 3D). These results can help interpret enigmatic skeletal effects of deletions of the presenilins and the Notch ligand Jagged at early stages of limb development (Pan et al., 2005) (see Conclusion).

### An empirically based mathematical model predicts the role of Hes1 synchrony in temporally coordinating condensation formation via effect on Gal-1A and Gal-8 levels

Our observation of Hes1 cyclic expression led us to hypothesize that the synchronization of cell fates brought about through cell-cell interaction (e.g., the juxtacrine signaling mediated by Notch and its ligands) was important for the temporal uniformity in emergence of condensations in vitro and in vivo. We first tested this hypothesis computationally. We modified a mathematical model that had been constructed to represent the behavior of limb mesenchymal cells via their biosynthesis and adhesive/de-adhesive interactions with Gal-1A and Gal-8. Previously, this model was shown to accurately simulate, in a one-dimensional interval, the morphogenesis of condensations and the conditions of their formation on the basis of a reaction-diffusion-adhesion pattern-forming instability (Glimm et al., 2014). Here we have introduced an additional function to the model *ϕ*(*x,t*), representing the phase of the Hes1 cycle, and made the cell response to the adhesive function of Gal-1A (“adhesion flux”) depend on it (see further description of the model in Materials and methods). This extended model was devised in analogy to the hypothesized clock-and-wavefront mechanism of somitogenesis, where the epithelization of somatic plate mesoderm occurs only during a specific phase range of the Hes1 cycle (Palmeirim et al., 1997; (Aulehla and Pourquie, 2006; Chal et al., 2017).

We considered four scenarios: (1) oscillations of the Hes1 molecular clock were in phase across the field of cells (‘oscillatory synchronous’); (2) the initial phase distribution was random, but cells oscillated with the same frequency (‘oscillatory asynchronous’);(3) Hes1 did not oscillate and its expression levels from cell to cell were random (‘non-oscillatory random’); and; and (4) there was no oscillation in Hes1 expression but it increased or decreased exponentially in time (‘non-oscillatory monotonic’). This simulates non-oscillatory decreasing or increasing Hes1 production rates. We simulated the temporal emergence of cell density spatial distribution in all four of these cases (Figs. 4).

**Figure 4:**
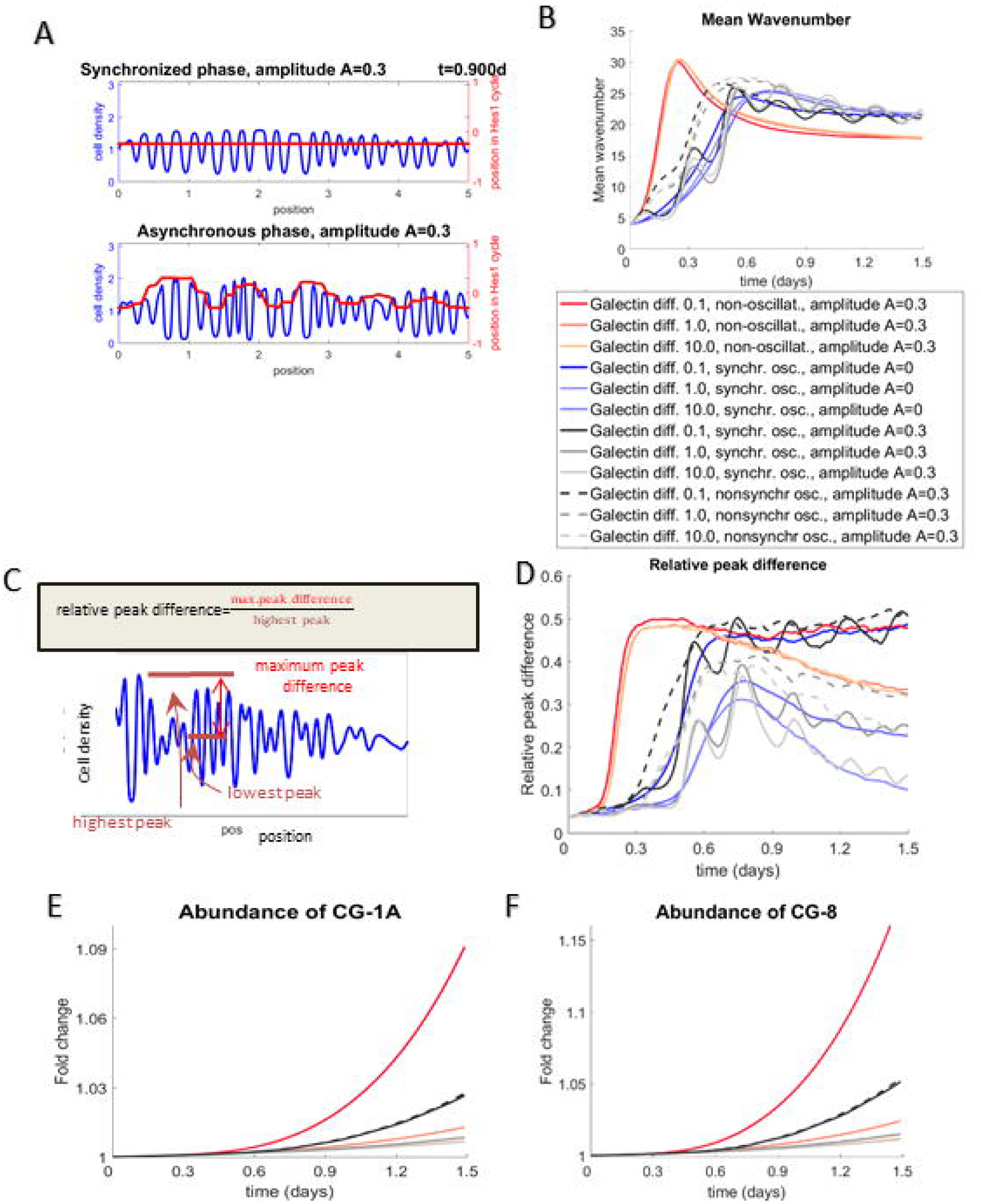
A model integrating oscillation in cell state with a reaction-diffusion-adhesion-based patterning mechanism predicts a role for synchronization in the spatiotemporal coordination of condensations. (A) Sample simulation. Top: Cell density of synchronous phase (blue) and state in the Hes1cycle. (The latter is sin*ϕ*, where *ϕ* is the Hes1 phase.) (red). Bottom: Cell density and state in the Hes1 cycle for asynchronous phase. Note that the bottom cell density appears less regular. (B) Fourier wavenumber of the patterns as a function of time. Note that there is very little difference between the synchronous and the asynchronous models, meaning that the typical length scale is unaffected by (lack of) synchrony. The wave number is reduced in the non-oscillating case of spatially random, but temporally fixed Hes1. (C) Formula for ‘relative maximum peak difference’, a parameter that measures the regularity of condensation patterns (see also Materials and Methods section titled ‘Simulations’) (D) Relative maximum peak difference (defined in Figure 2.1 B) as a function of time based on N=75 simulations of synchronized Hes1 phases (solid line) and asynchronous Hes1 phases (dashed) for four different values of the galectin diffusion coefficient d_*gal*_ = 0.1,0.5,1,10 as a fraction of the standard value 11.5 μm^2^/s. The relative amplitude of the effect of the Hes1 oscillations on the cell-cell adhesion flux coefficient was *A=0*.3 except for the case with constant zero amplitude A=0. (E,F) Predicted abundance of Gal-1A and −8 over time gives a mild increase in Gal-1A and Gal-8 abundance when Hes1 phases are synchronous relative to the condition when the phases are asynchronous. This difference is only observed when galectin diffusion coefficient d_*gal*_ = 0.1. Key (bottom) shows the temporal dynamics of parameters measured in B, C and D at different galectin diffusion coefficients and when phases are synchronous (solid lines) versus asynchronous (dashed lines). Both galectins increase at much higher rate in the non-oscillating Hes1 case (red lines).

We observed that in both the ‘oscillatory synchronous’ and ‘oscillatory asynchronous’ cases spatial patterns emerged (Fig. 4A and Movie S2), which essentially had the same Fourier wavenumber, i.e., the number of condensations per unit length was essentially unaffected by synchrony of the Hes1 oscillations (Figure 4B). The wavenumber was slightly lower in the case of ‘non-oscillatory random’ Hes1 (Figure 4B). However, patterns that emerged in the ‘oscillatory synchronous’ case were markedly more regular than the patterns of the ‘oscillatory asynchronous’ counterpart. For ‘non-oscillatory random’ Hes1, this irregularity was even more pronounced (Movie S3). The regularity of patterns was quantified via a new measure, termed the ‘relative maximum peak difference’ (Fig. 4C). This is the difference in density between the densest condensation and the least dense condensation, i.e., highest and the lowest peak in the cell density, relative to the highest density. This quantifies the uniformity of the pattern density across the different aggregation sites, with values close to zero indicate very regular patterns. The spatial regularity of the amplitude of cell density distribution was reproducibly diminished in the ‘oscillatory asynchronous phase’ case. This difference was even more striking in the case of ‘non-oscillatory random’ Hes1. Whereas in both the oscillatory synchronous and oscillatory asynchronous phase cases, the irregularity tended to decrease with time through coalescence and re-arrangement of the cell density, this effect was hardly present in the case of non-oscillating random Hes1 (Fig. 4D).

Another measure of pattern regularity is the variance of the Fourier wavenumber of the cell density pattern, with smaller standard deviations indicating more regular patterns (e.g., a perfect sine wave having zero standard deviation). Here too, the standard deviation was slightly lower in the synchronized case, indicating greater regularity of those patterns (Fig. S3). In fact, synchronizing the intracellular oscillatory phases after starting the simulations with random spatially correlated phases ‘rescued’ the uniformity in condensation patterning as well (Movie S4). While the standard deviation of the wavenumber was slightly higher in the ‘oscillating asynchronous’ Hes1 case than in the ‘oscillating synchronous’ Hes1 case, indicating less regular patterns, it was much higher in the ‘non-oscillating random’ Hes1 case, thus giving more quantifiable evidence that this case produces the least regular patterns.

We also performed simulations in which the amplitude A of the term describing the effect of oscillations of the Hes1 state was set to zero (Fig 4). This corresponds to the case of effectively non-oscillatory, but spatially homogeneous Hes1 expression. In other words, the Hes1 state is constant in both time and space. The resulting patterns are very similar to the case of nonzero amplitude A (Fig 4, where the comparison is with A=0.3), which is the oscillatory synchronous Hes1 scenario. The mean wave number, the standard deviation of the wavenumber and the relative peak difference of the A=0 case are all almost indistinguishable from the temporal averages of the oscillatory synchronous Hes1 scenario (Fig 4 and S3). These comparisons, along with the irregularity of the oscillator asynchronous Hes1 scenario, indicate that it is not so much the oscillatory nature of Hes1 expression which is crucial for pattern regularity, but rather the spatial uniformity of the Hes1 state. The importance of synchronous oscillations may thus be as a device for ensuring this uniformity of Hes1 state.

One crucial parameter was the diffusion coefficient of the two galectins, which were assumed to diffuse at the same rate. The smaller the diffusion coefficient, the more irregular are the induced cell density patterns (Fig. 4D). We found that in the ‘oscillating asynchronous’ Hes1 case, the overall expression rates of the genes specifying both Gal-1A and Gal-8 were predicted to be mildly elevated compared to the ‘oscillating synchronous’ Hes1 case (Figs. 4E and 4F). The effect was more pronounced in simulations for which the galectin diffusion coefficients were smaller. This was much more pronounced in the case of ‘non-oscillating random’ Hes1, where both Gal-1A and Gal-8 increased at a much higher rate (Figs. 4E and 4F).

The simulations of the increasing or decreasing non-oscillatory Hes1 case showed the following results: In the case of *decreasing* Hes1 production, the wavenumber increased compared to the control case of spatially synchronous oscillating Hes1 production with a constant temporal mean (the synchronous case). Thus, there were more condensations per unit length (Fig 4A). For very large wavenumbers, the resulting patterns of densely distributed condensation may be interpreted as continuous endoskeletal plates, like those seen in cartilaginous and ray-finned fish (Bhat et al., 2016). Also, the faster the decrease in Hes1 production, the faster was the rise of the overall expression rates of Gal-1A and Gal-8 (Fig. 4B and C). We also simulated another scenario, where Hes1 *increased* with variable rates without oscillating. We observed extremely low wave number (prognosing the absence of pattern by the criteria considered here) across diverse diffusivities of Gal-1A and Gal-8 (Fig 4A). With increased rates of Hes1 expression, the levels of Gal-1A and Gal-8 were predicted to decrease below control values (Fig. 4B and C). The simulation results indicate that both potential effects – loss of synchrony of Hes1 oscillations and non-oscillatory change of Hes1 expression rate – lead to less regular condensation patterns, with a greater impact of the former than the latter. The two effects may occur together, such as under the influence of a γ-secretase inhibitor such as DAPT which results in an attenuation of Hes1 expression (Fig. 3A).

An assumption of our oscillation-enhanced mathematical model is that the galectin-based condensation patterning network is under the regulation of synchrony-inducing cell-cell communication. As predicted by the model (Figs. 4E and F), the messenger RNA levels of genes coding for both Gal-1A and Gal-8 were found to be significantly elevated in cultures pretreated with DAPT (Fig. 5). Indirect immunocytochemistry showed that DAPT treatment also brought about an increase in Gal-1A protein staining (Fig. S4)

**Figure 5:**
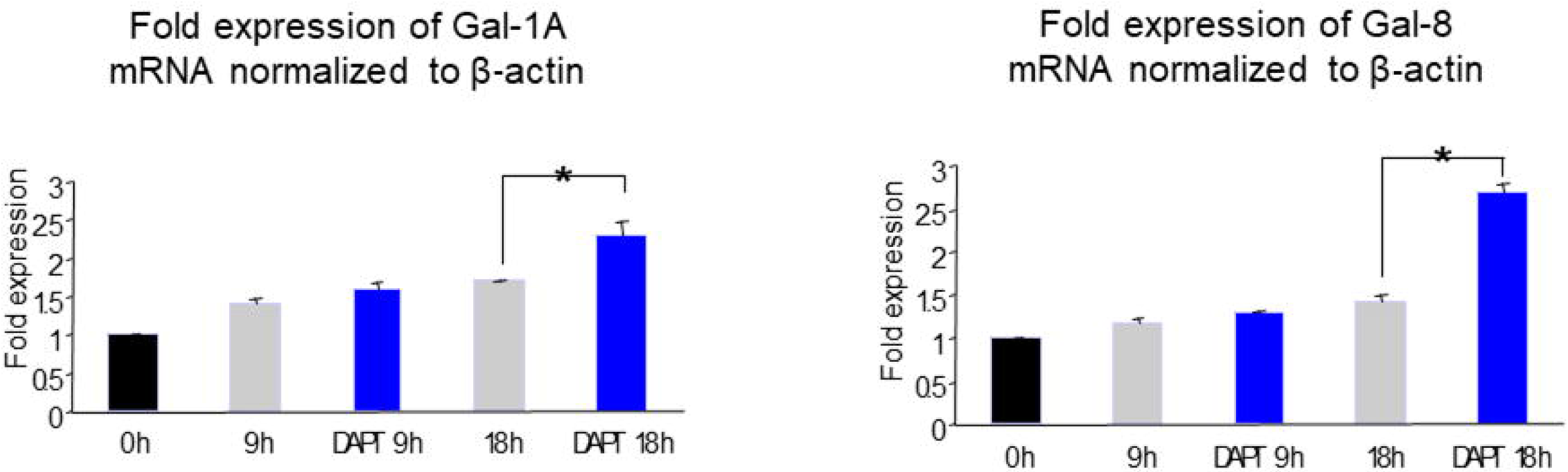
The γ-secretase inhibitor DAPT enhances the gene expression of Gal-1A and Gal-8. Treatment with DAPT increased expression of Gal-1A (left) and Gal-8 (right) when assayed relative to untreated micromasses at 9 hours and 18 hours post culture initiation. Error bars represent S.E.M. *p < 0.05.

### Other treatments that block Hes1 oscillation alter Gal-1A levels and condensation patterns

In earlier reports, TGF-β signaling was shown to alter the patterning of limb mesenchymal condensations (Leonard et al., 1991). Treatment with TGF-β brought about condensations which were larger and fused. Given its positively autoregulatory role, TGF-β had been hypothesized to be the ‘activator’ morphogen in a reaction-diffusion-based mechanism of limb skeletal patterning (Hentschel et al., 2004). Consistent with this hypothesis, mesenchyme-specific knockout of the type II TGF-β receptor Tgfbr2 led to (depending on the context) both positive and negative effects on limb chondrogenesis (Seo and Serra, 2007). We therefore treated limb precartilage micromasses with SB505124, a small-molecule TGF-β receptor antagonist, and elicited a decrease in mean size of the condensations (Fig. 6 top panel). This was concomitant with a decrease in micromass staining for Gal-1A by indirect immunocytochemistry (Fig. 6 middle panel, see also Fig. S5). We therefore looked at the effect of TGF-β signaling inhibition on *hes1* oscillations, the modulation of which had marked effects on condensation patterning (Fig. 6). When Hes1 mRNA was assayed at diverse time points, it was found to be consistently upregulated without any oscillatory dynamics (Fig. 6 bottom row). Therefore, the inhibition of TGF-β signaling impaired condensation dynamics in association with both decreased Gal-1A levels and a non-oscillatory elevated expression state of *hes1.*

**Figure 6:**
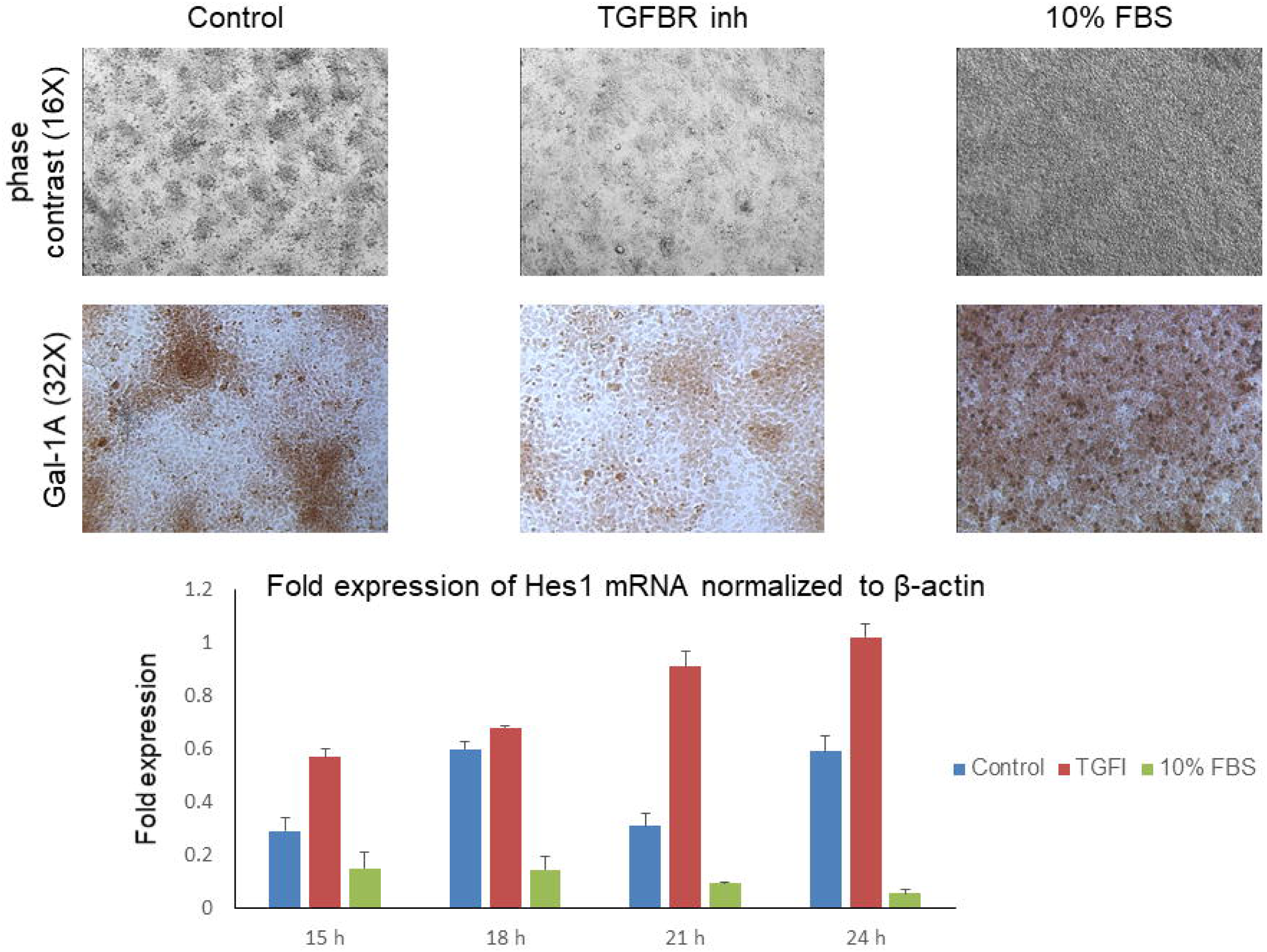
Exogenous treatment with TGF-β signaling inhibitor and serum had opposite effects on condensation pattern, Gal-1A levels and Hes1 expression. Treatment of micromasses with SB505124 (5 μM) resulted in smaller and more widely spaced condensations with an overall increase in the number of cells that did not stain for Gal-1A. In contrast, exposure to FBS led to formation of a single large condensation, with all the cells within it showing high levels of Gal-1A. The SB505124 treatment caused a sustained non-oscillatory increase in Hes1 levels, whereas serum exposure had the opposite effect: there was a sustained non-oscillatory decrease in Hes1. Error bars represent S.E.M. *p < 0.05.

This result raised the question of whether the effect of impairment of *hes1* oscillations on chondrogenic patterning was solely due to an elevation of Gal-1A expression. This possibility was disconfirmed by the results of treatment of cultures with fetal bovine serum, an inducer of chondrogenic differentiation and of recruitment of intercondensation mesenchyme into cartilage nodules (Downie and Newman, 1994). Serum treatment resulted in an increase in Gal-1A levels and sustained (i.e., non-oscillatory) low levels of Hes1 mRNA (Fig. 6). Our experiments indicate that treatments that decreased or increased Hes1 expression had predictably opposite effects on the patterning of condensations, consistent with our results above, and our model, for the involvement of this transcriptional regulator in the process. The treatments also stopped the oscillation. While this may be a side-effect of the treatments with no definite implication for the oscillator’s role (although TGF-β signaling directly regulates *hes1* transcription (Peignon et al., 2011), one interpretation of these results is that nearby cells need to have similar Hes1 levels in order to respond similarly to similar levels of an inductive signal (e.g., Gal-1A). Having Hes1 oscillate, and having the oscillation synchronize, ensures that noisiness in this factor will not degrade the pattern.

## Discussion

The concept of the morphogenetic field has been intermittently influential in developmental biology for close to a century (reviewed in (Gilbert et al., 1996) and Beloussov et al. (1997)). Joseph Needham, using a definition he attributed to C.H. Waddington, characterized a morphogenetic field as “a system of order such that the positions taken up by unstable entities in one part of the system bear a definite relation to the position taken up by other unstable entities in other parts of the system” (Needham (1937) p. 71). To biologists in the modern molecular and physics-informed period, the global coordination described has been variously attributed to competing signaling centers, the transcriptional effects of which are regulated by different extracellular morphogens (Zakin and De Robertis, 2010), electrical fields, in turn generated by and promoting, ion fluxes across cell boundaries (Levin, 2012), and quorum sensing in cell collectives mediated by morphogens and other transported signals (Widelitz and Chuong, 2016).

Our experimental observations, in combination with the oscillation-enabled version of the theoretical model of Glimm et al. (2014) presented in this paper strongly suggest that the synchronization of cell fates through γ-secretase-Hes1-dependent signaling results in spatiotemporal coordination of the patterning of digital condensations in micromasses in vitro and in the digital plate in situ. Synchronization of cellular oscillators does not require transport of materials like morphogens or ions between or through a field of cells and is therefore an apt mechanism for the global coordination implied by classical notions of the morphogenetic field. The lack of dependence on molecular transport means that it can occur rapidly over linear distances of tens to hundreds of cells, in suspensions or tissues (Chen et al., 2017; Garcia-Ojalvo et al., 2004; Ozbudak and Lewis, 2008). Synchronization can result from weak non-specific coupling between neighboring subunits that oscillate autonomously (Strogatz, 2000) or alternatively may be a consequence of global coordination of oscillations that only emerge when cells interact with one another via juxtacrine-type mechanisms (Hubaud et al., 2017). The latter include transmembrane ligand-receptor combinations as Notch-Delta, but also extracellular ligand-receptor binding that brings cells in close apposition with each other (Gilbert, 2010).

Where oscillations of the gene regulatory factor Hes1 are synchronized, as occurs in the presegmental plate mesoderm of vertebrate embryos (Giudicelli et al., 2007; Ozbudak and Lewis, 2008) and in the digital plate mesenchyme described here, the levels of this transcriptional co-regulator are rendered identical across a developing primordium at each phase of development. Therefore, our results are consistent with the idea that it is not oscillation per se that is important for the coordination and refinement of condensation formation, but uniform Hes1 status across the tissue field. Synchronized oscillations are an efficient way, using available means, to achieve this across long distances relative to the scale of single cells. Artificial treatments can force this condition in the absence of oscillation. Thus, in mouse limb micromass cultures, the effect of DAPT in causing fusion of precartilage condensations (as reported here for chicken; Fig. 5A), was counteracted by exogenous Notch1 intracellular domain (NICD) (Fujimaki et al., 2006).

Hes1 spatial uniformity potentially eliminates a major source of randomness in responses of distant cells to patterns of morphogenetic factors generated by resident mechanisms (e.g., diffusion gradients; reaction-diffusion processes). The oscillation synchronization mechanism also ensures that Hes1 values, however they vary over time, will reside within a “normal” range, defined by the extrema of the oscillation, unlike the “abnormal” values that may be induced by Hes1 perturbants such as TGF-β or serum (Fig. 6). This global coordination phenomenon, evidenced in the remarkable simultaneity of emergence of condensations across a micromass culture (Movie S1), precisely fulfils the Needham-Waddington criterion of “unstable entities” in one part of the system being correlated with such entities in other parts of the system.

In an earlier study, *c-hairy 2*, a paralog of *hes1*, was shown to cycle with a 6 h periodicity in developing chicken limb autopodial mesenchyme (Pascoal et al., 2007), suggesting that the cyclic gene expression dynamics we have observed may have multiple components. Neither our study nor the earlier one tested the direct involvement of Notch and its ligands on mediating synchronization of Hes gene expression. However, the dependence on γ-secretase reported here, as well as the expression of Notch1 in the chicken digital plate at the appropriate stages (Williams et al., 2009) make a strong circumstantial case for a role for this pathway, as do the results of Fujimaki et al. (2006) employing forced expression of NICD, cited above. As a caveat, however, we note that DAPT also inhibits steps in the TGF-β pathway (Pazos et al., 2017), and that the responsiveness of limb bud mesenchyme to TGF-β with respect to expression of several chondrogenesis-related parameters is also cyclical (Leonard et al., 1991). Therefore, like the mesenchymal clock itself, the medium of its synchronization may be multifactorial.

If, as we suggest, the responsiveness of precartilage mesenchymal cells to elevated levels of Gal-1A depends on the cells’ Hes1 status, synchronizing *hes1* gene expression across the tissue ensures that local variations of on-off responses in the digital plate will be sharp rather than noisy, leading to well-formed skeletal elements with smooth boundaries. In vitro, in the absence of nonuniformities such as Shh and Hox protein gradients, this synchrony has the effect of making all the condensations appear simultaneously.

Our results are consistent with, and help interpret, earlier genetic experiments in which Presenilin 1 and 2 or other components of the Notch signaling pathway were manipulated in the developing limb ectoderm or mesenchyme. The loss in the apical ectodermal ridge of both presenilins led to fusions and truncations of digits, with variable penetrance (Pan et al., 2005). Based on our experiments and computational simulations this would be a predicted result of loss of uniform Hes1 status across the digital plate. In our cultures, residual ectodermal cells could ‘seed’ Hes1 expression in the mesenchymal micromass or digital plate, which would then become self-sustaining through Notch-based or Notch-independent γ-secretase regulated signaling. Similarly, self-sustained synchronization of Hes1 expression may result from early exposure to ectoderm in the experiments of (Pan et al., 2005), accounting for what these investigators characterize as a mesenchymal “memory” of the ectoderm-dependent Notch signaling that occurs during early development and is necessary for reliable chondrogenic patterning.

## Materials and methods

### Chicken eggs

White Leghorn chicken fertilized eggs were obtained from Moyer’s Chicks, Quakertown, PA. Eggs were incubated in a humidified incubator at 39°C for 5-8 days.

### Micromass cell culture

Primary high density micromass cultures were performed as described previously (Bhat et al., 2011). Briefly, mesenchymal tissue was dissected from the myoblast-free autopod field of leg and wing buds of stage 24 (Hamburger and Hamilton, 1951) chicken embryos (∼4½ d of incubation), dissociated with trypLE Express solution (Gibco), filtered through Nytex 20-*µ* m monofilament nylon mesh (Tetko, Briarcliff Manor, NY) and resuspended in medium for plating at 2.5×10^5^ cells per 10 *µ* l spot. Cell spots were deposited in Costar 24 well tissue culture plates and allowed to attach for 1 h before the wells were flooded with 1 ml of serum-free defined medium (henceforth called DM, (Paulsen and Solursh, 1988)): 60% Hams F12, 40% Dulbecco’s modified Earle’s Medium (DMEM), 5 μg/ml insulin, 100 nM hydrocortisone, 50 *µ* g/ml L-ascorbic acid, 5 *µ* g/ml chicken transferrin). Medium was changed daily.

In experiments to assay for effects of Notch signaling on galectins or synchronization of condensations, cultures were treated for specified time periods with 500 nM-5 *μ* M DAPT (N- [N- (3,5-difluorophenacetyl-L-alanyl)]-S-phenylglycine t-butyl ester) (Sigma Aldrich). For serum treatment, cells were cultured in DM supplemented with 10% fetal bovine serum. For inhibition of TGF-β signaling, cells were treated with 5 μM SB505124, an ALK4/5/7 antagonist (R&D systems, Minneapolis, MN).

### Quantitative PCR

Total RNA was isolated from cells according to instructions included with the Absolutely RNA Microprep kit (Agilent) as described previously (Bhat et al., 2011). RNA isolated from cells from micromass cultures grown to specified time points or freshly dissected and dissociated 5 d limb buds was quantitated with a Nanodrop ND-1000 spectrophotometer at 260 nM and used as template for the synthesis of cDNA. Comparative quantitative PCR (qPCR) was performed on the Mx3005P instrument (Stratagene) with Brilliant SYBR Green QPCR master mix (Stratagene). The amplification protocol for comparative quantitative PCR included the following steps: denaturation at 95°C for 10 min, 40 amplification cycles (95°C for 30 sec, 55°C for 1 min, 72°C for 1 min) and melting the amplified DNA in order to obtain melting curve for the PCR product. Messenger RNA expression of the gene of interest was normalized to internal control gene β-actin and analyzed by the ΔΔCt method (Livak and Schmittgen, 2001). At least two biological replicates were prepared for all analyses.

### Whole mount *in situ* hybridization

In situ hybridization was performed on whole embryos essentially as described (Acloque et al., 2008). cDNA for chicken Hes1 gene (kind gift from Dr. Olivier Pourquié) was inserted into pGEM-4Z vector (Promega) and linearized with *Eco*RI for synthesis of labeled and antisense strands using SP6 polymerase and the digoxygenin-based DIG RNA labeling kit (Roche), or with *Hin*dIII for synthesis of the labeled sense strand using T7 polymerase and the DIG RNA labeling kit (Roche). All the prehybridization and posthybridization washes were performed using the BioLane HTI 16V tissue processing robot (Intavis Bioanytical Instruments, Koeln, Germany).

### Indirect immunostaining

Limb mesenchymal cultures were fixed with 100% methanol precooled to −20°C for 10 min. After three washes with PBS for 5 min each, the fixed cultures were blocked with 10% non-immune goat serum (Histostain-Plus kit; Zymed). and incubated for 1 h with 5 μg/ml polyclonal rabbit anti-CG-1A, or 5 μg/ml polyclonal rabbit anti-Hes1 antibody (AB-15470, Millipore; Antibody Registry # AB_11213583). Following the Histostain-Plus protocol, the cultures were incubated for 10 min each with a biotinylated broad-spectrum secondary antibody, streptavidin peroxidase conjugate and a 1:2:1 combination of 0.6% hydrogen peroxide, diaminobenzidine (DAB) substrate solution and a substrate buffer. Rinsed cultures were viewed through a Zeiss IM 35 inverted microscope at 16x or 32x objective magnification and a Zeiss binocular dissecting microscope at 1.2 x magnification to discern the staining pattern of the whole micromass culture. Controls were incubated without primary antibodies, or secondary antibodies. Quantification of the cytochemical staining was performed using IHC Profiler Macro plugin (Varghese et al., 2014) for the ImageJ software package (http://www.rsb.info.nih.gov/ij/) (Schneider et al., 2012).

### In ovo manipulation of wing buds

Chick embryos were incubated horizontally. On the third day, the eggs were candled to locate the position of the embryos and 3-4 ml of albumin was removed by puncturing the wide bottom of the egg with an 18-gauge needle attached to a syringe. On day 4 the shell overlying the embryo was carefully removed and the window in the egg sealed with sterile transparent tape. Between 5-6 days ∼1 *µ* g of DAPT (2 *µ* g/ml in nuclease-free PBS) was injected into the mesenchyme of the autopod field of the wing bud. Control embryos were injected with identical amounts of BSA. The egg was resealed and the embryos were allowed to grow for 2-3 more days.

### Whole-mount skeletal preparation

Embryos at 8-9 d were washed in PBS and fixed in 5% trichloroactetic acid overnight. Embryos were stained 0.1% Alcian blue in acid alcohol overnight. Following destaining in acid alcohol overnight, embryos were dehydrated in absolute ethyl alcohol and cleared in methyl salicylate.

### Statistics of pattern formation

Experiments that involved the inhibition of protein function and pharmacological treatment appeared to bring about a change in pattern formation. To quantitatively assess the effect of these agents on the condensation pattern, three parameters were assessed separately. These were condensation size, condensation number and intercondensation distance. The cultures were photographed under low magnification (2.5×) and the images were binarized using ImageJ. The smallest condensation was chosen by eye from among all the culture pictures and was set as the threshold for the lower limit of condensation size measurement. Image J was then used to automatically measure the number and sizes of condensations in each picture.

To measure inter-condensation distances, an arbitrary point was chosen on the boundary of each condensation. This point was then connected with the point on the boundary of the condensation that was farthest from it, such that the line connecting the two points lay within the condensation. Since most of the condensations were quasi-circular, the line joining the two points represented an approximation to the diameter of the condensation. The mid-point of this axis was taken to be the center of the condensation. The centers of the condensations were then connected to those of neighboring condensations and the intercondensation distances were measured.

The values of the condensation numbers, sizes and inter-condensation distances were represented as fold means +/-SEM. The difference between groups and controls was assessed by two-tailed unpaired parametric Student’s t-test and taken to be significant if the p-value was <0.05. Experiments was performed 3 or more times with 3 or more technical replicates unless otherwise specified; additionally, each micromass culture contained randomized cells derived from the distal portions of 20-30 embryonic chick wing or leg buds.

### The Lomb-Scargle algorithm

The Lomb Scargle algorithm is a statistical tool used to search for periodicity of fluctuating data when the data sets are incomplete or when the data collection is not stringently distributed over the entire period of data collection (Glynn et al., 2006). The algorithm constructs a periodogram (a graph in which peaks indicate frequencies with significant periodicities) with known statistical properties. The L-S algorithm has advantages over other methods like Fast Fourier Transforms (FFTs) which require evenly sampled data, and has been used to identify oscillating genes involved in vertebrate segmentation (Dequeant et al., 2006).

### Mathematical model

We used the following system of partial differential equations, which were adapted from Glimm et al. (2014), equations (4.2)-(4.4):

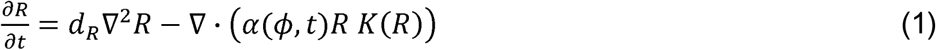

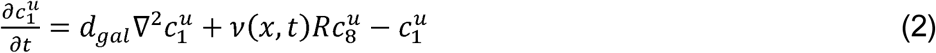

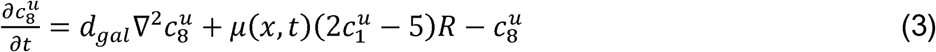

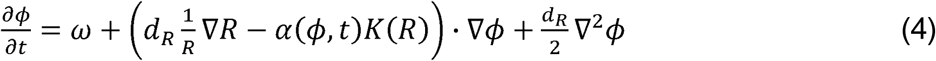

Here *R*(*x,t*) is the cell density and 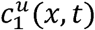 and 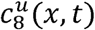 are the concentrations of diffusible CG-1A and CG-8, respectively, as functions of time *t* and position *x*. The intracellular Hes1 oscillations are modeled by an effective mean oscillation phase *ϕ* (*x,t*). The above equations model the diffusion and expression of galectins, as well as random motion of cells and cell-cell adhesion. The diffusion coefficient of the cell density was *d*_*R*_ = 0.08, while the galectin diffusion coefficient *d*_*gal*_ was varied around the standard value of *d*_*gal*_ = 1. The latter is modeled through an effective adhesion flux term (following Armstrong et al, 2006) given by

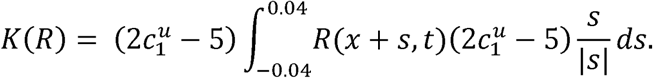

To account for the oscillatory dynamics of Hes1, we consider the phase of the Hes1 cycle *ϕ* (*x,t*) as one of the variables. Its spatiotemporal evolution is governed by a constant rate of change *ω*, a small diffusion term to account for local interactions, and an advection term that models the effective transport of the phase with the cells (Jörg et al., 2016). The constant temporal rate of change *ω* was chosen as 8*π*, corresponding to a period of 6 hours.

Crucially, the effect of the state of Hes1 in its oscillatory cycle on the galectin genetic regulatory network was modeled via making the cell-cell adhesion flux dependent on the Hes1 phase *ϕ* (*x,t*). This is a ‘high level’ modeling approach: Since the details of the interactions are unknown, we incorporate their ultimate effect on cell movement rather than directly implementing their direct molecular effects. Specifically, we set

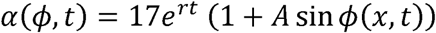

where 0 ≤ *A* ≤ 1 is a parameter that describes the relative amplitude of the oscillation. The parameter *r* encodes the effect of increasing or decreasing Hes1 base production; it is the temporal rate of exponential growth (for positive *r*) or decay (for negative for *r*).

### Comparison to previous model

Our original model (Glimm et al., 2014) did not incorporate the Hes1 phase *ϕ* (*x,t*). There the main concern was the analysis of the galectin regulatory network and how, and under what conditions it undergoes spatial symmetry breaking to form patterns of precartilage condensations. The present paper deals with a second order problem: the control of the regularity of patterns in space and time. Apart from incorporating an expression representing the experimentally inferred Hes1 oscillations, the present analysis also required more attention to the exact form of the initial conditions.

The simultaneous emergence of density peaks in Glimm et al. (2014) were a consequence of choosing initial conditions as random, spatially uncorrelated perturbations of a spatially homogeneous steady state of the cell density. The resulting initial conditions lacked a characteristic spatial scale, making them effectively uniform, which in turn led to a highly synchronous and uniform patterning. In the paper at hand, we use random initial conditions for both the cell density and the Hes1 phase, which are spatially correlated. (For the details of how the initial conditions were set up, see the next section.) This adds biological realism, since it means that sites neighboring foci of elevated cell density have an increased likelihood of higher cell density as well. (For the details of how the initial conditions were set up, see the next section.) As a result, this yielded less regular condensation patterns in the absence of a global coordinating mechanism, as expected in any extended system employing only local interactions.

### Simulations

In the first set of numerical simulations (‘synchronized phase’), the initial phase *ϕ* (*x,*0) was chosen as the spatially constant value 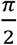, thus giving the spatially uniform phase 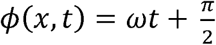. In the second set of simulations (‘asynchronous phase’), a random distribution for the initial phase *ϕ* (*x,*0) was generated as follows: Using a spatial discretization of a one dimensional domain with N=1000 evenly spaced grid points, we first defined an ‘influence function’ *I*_*n*_ (for integer values *n*) such that *I*_*n*_ = 1 for −4 <*n*< 4, *I*_*n*_ = 0 for *n* < −303 or *n* > 303, a linear function with slope 1/300 for −303 ≤ *n* ≤ −4, interpolating between 0 and 1, and similarly a linear function of slope −1/300 for 4 ≤*n* ≤ 303, interpolating between 1 and 0. Starting with *ϕ* (*x*) = π/2, we then choose a random point *n*\* and a number p which is −1 or +1 with equal probability and modify *ϕ* (*x*) by adding 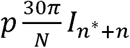 to it, where this expression is to be understood cyclically. This process is repeated 15*N* times to generate a random phase *ϕ* (*x*) with mean π/2. The system procedure was applied to produce the initial cell density *R*(*x*), with the only differences being that the starting value was *R*(*x*) = 1 and the addition in each step was 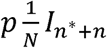, using the same notation as above.

To simulate the ‘non-oscillating random’ Hes1 case, a random phase *ϕ* (*x*) with mean 0 was generated with the procedure described above. However, this phase was kept temporally constant by replacing equation (4) by the equation 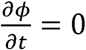.

The system was solved on the one-dimensional interval [0,5] with periodic boundary conditions using a spatial discretization with N=2500 evenly spaced grid points and the method of lines with Matlab’s ode45 algorithm, a variable step Runge-Kutta ODE solver. The resulting final cell densities were analyzed using the Discrete Fourier Transform. We also used a new measure we call the relative maximum peak difference, which is the difference between the largest local maximum and the smallest local maximum as a fraction of the largest local maximum. This is a measure of the regularity of the pattern; the larger this maximum peak difference, the less regular we may consider the pattern to be. We generated statistics with n=50 or n=75 runs of each set of simulations.

### Movie Links

Movie S1: https://www.dropbox.com/s/oon1s8nfjkz57n3/Movie%20S1.avi?dl=0

Movie S2: https://www.dropbox.com/s/k3ixk3qu3nl6560/Movie%20S2.mp4?dl=0

Movie S3: https://www.dropbox.com/s/587rde5cvpgvgsp/Movie%20S3.mp4?dl=0

**Table 1.**
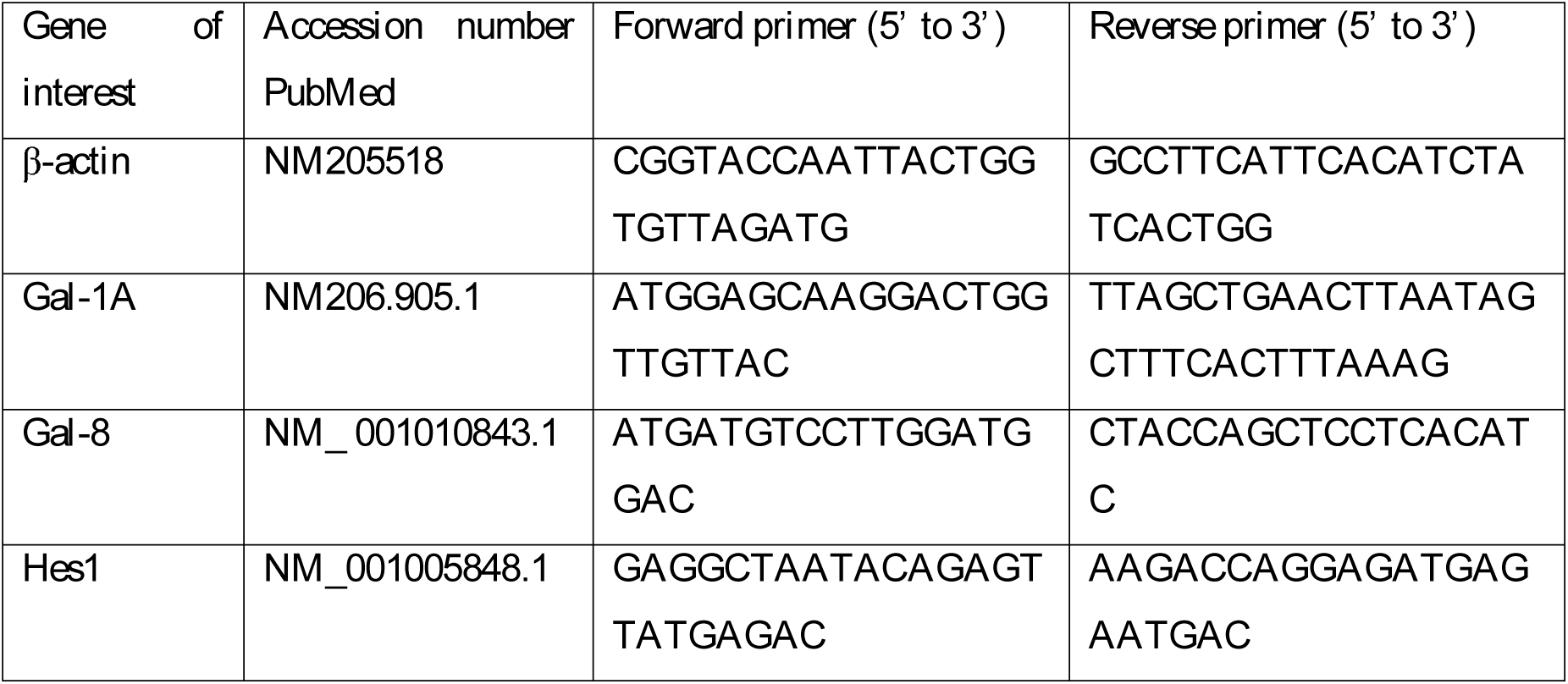
Primer sets used for qRT-PCR.

---

**Movie S1:** Time-lapse video of a high-density leg bud micromass culture photographed using fluorescence optics. The cell layer, which is confluent from the start, contains a subpopulation (approximately 5% of the total) that were electroporated with a DNA plasmid directing the expression of H2B-GFP. Precartilage condensations emerge in a synchronous fashion over the entire 2.5 mm diameter culture (the field of view in the video is about 2 mm wide) becoming clearly visible by the end of two days and forming cartilage nodules over the next day and a half.

**Movie S2:** Comparison of two simulations of cell condensation in one spatial dimension using the mathematical model described in the “Materials and Methods” section. The simulated time is between 0 days and 1.5 days. The blue curve shows the cell density (Rt,x) as a function of space, normalized so that the initial uniform density is 1. The red curve shows the fraction 0.3 sinφt,x of the sine of the Hes1 phase. Top panel: Oscillatory synchronized Hes1 phases. Bottom panel: Oscillatory asynchronous phases. The random initial cell density is the same for both panels. The galectin diffusion coefficient is 1. The condensation pattern emerges more synchronously in the synchronized phase case and the resulting pattern is more uniform. (Other parameters: A= 0.3)

**Movie S3:** Comparison of two simulations of cell condensation in one spatial dimension using the mathematical model described in the “Materials and Methods” section. The movies show two simulations of the “non-oscillatory random” Hes1 scenario. The two cases have the same initial conditions and the parameter sets differ only in the galectin diffusion coefficient (0.1 on top and 10 on the bottom). The presentation is the same as in movie S2, but note that the blue line representing the Hes1 state does not oscillate. Note how both final patterns are irregular in that there is no obvious characteristic wavelength, and this lack is slightly more pronounced in the case of the lower diffusion coefficient.

**Movie S4:** Comparison of two simulations of cell condensation in one spatial dimension using the mathematical model described in the “Materials and Methods” section. Parameters and presentation is the same as in movie S2. The initial simulated time period of 1.5 days is the same for both panels which show random oscillation. At that point in time, the phase φ(t,x) is spatially synchronized in the bottom (‘rescue’) panel. The condensation pattern emerges more synchronously in this ‘rescued’ panel and the resulting pattern is more uniform.

